# Rapid early life colonization of the intestinal tract by *Akkermansia muciniphila* after voluntary feeding

**DOI:** 10.64898/2026.02.06.704081

**Authors:** Jenice M. Dumlao, Paige McCallum, Colten R. Hodak, Elizabeth Guinto, Winnie Enns, Lauren Davey, Jonathan C. Choy

**Affiliations:** Department of Molecular Biology and Biochemistry, Simon Fraser University, Burnaby, British Columbia, Canada; Department of Biochemistry and Microbiology, University of Victoria, Victoria, British Columbia, Canada

**Author notes:** <u>Corresponding author</u>: Jonathan Choy, PhD, Department of Molecular Biology and Biochemistry, Simon Fraser University, Burnaby, British Columbia, Canada.

**Keywords:** Gut microbiota, Akkermansia muciniphila, Dysbiosis, Oral administration, Probiotics

## Abstract

**Background:** Non-invasive methods to colonize intact gut microbiota populations with specific bacterial species are useful for experimental studies that advance our understanding of this commensal microbial population in health and disease. Within the gut microbiota, the anaerobic muciniphile *Akkermansia muciniphila* has many established health benefits. We report the development of a new voluntary feeding protocol for non-invasive administration of bacteria into the intestine and use it to characterize the early life colonization of the intestinal tract by *A. muciniphila*.

**Results:** Mice were voluntarily fed a human strain of *A. muciniphila* (MucT/BAA-835) in the week after weaning, whereupon they consistently and rapidly ingested the bacterium. At this developmental period, conventionally housed mice were rapidly colonized by human *A. muciniphila* that persisted until at least 8 weeks of age. In mice that contained a dysbiotic gut microbiota that lack endogenous *A. muciniphila*, voluntary feeding with human *A. muciniphila* similarly led to rapid and stable colonization. Colonization was similar in female and male mice. Also, in conventionally housed mice there was incomplete colonization of the intestinal tract with endogenous *A. muciniphila* between 3 – 4 weeks of age, which enabled its competitive exclusion by human *A. muciniphila* that was orally delivered.

**Conclusions:** These findings establish a new and non-invasive approach for colonizing the intestinal tract with commensal microbes that provides information on the early life colonization of the gut microbiota with *A. muciniphila*.

## Background

The community of commensal microbes that inhabits the intestinal tract of mammals, termed the gut microbiota, is essential for health. These microbes contribute to digestion of intestinal contents, control the absorption of nutrients and production of metabolites into the body, and affects the development of several other physiological processes essential for homeostasis [1]. Dysfunctional alterations in the composition of the gut microbiota, referred to as dysbiosis, contributes to diseases such as asthma, allergy, cardiovascular disease, and inflammatory bowel disease [2-5]. Preclinical studies have also implicated the gut microbiota in affecting the development of many other diseases ranging from multiple sclerosis to autism and anxiety [6, 7].

The composition of the gut microbiota is diverse among the human population. The major phyla known to be components of this community are Firmicutes and Bacteroidetes, which represent 90% of the gut microbial population, as well as Actinobacteria, Proteobacteria, Fusobacteria and Verrucomicrobia [8]. However, the abundance of bacterial genera that are found to inhabit the intestinal tract of an individual are highly variable and dependent on diet, age, sex, environment, and disease [9]. Although there is diversity in its specific composition, there can be convergence in the metabolic functions of bacteria in the gut microbiota within and between mammalian species [10]. In the setting of this diversity, *Akkermansia* spp. are the only genera found in the gut microbiota that use glycans from host mucin as its primary energy source, and *Akkermansia muciniphila* is the dominant species found in the gut [11]. It is a strict anaerobe that is consistently present in humans, albeit in variable abundance, as well as in many other mammals including laboratory mice [12, 13].

*A. muciniphila* is an important component of the healthy human gut microbiota and a reduction in its abundance is associated with the development of obesity, metabolic syndrome, and inflammatory bowel disease [14-16]. Stimulation of TLR2 by the *A. muciniphila* outer membrane protein Amuc_1100 induces anti-inflammatory responses within the intestine but live bacterium may also promote immunotherapeutic efficacy for some cancers [17-19]. Also, the metabolism of mucin by this bacterium leads to the production of short chain fatty acids (SCFAs) that have pro-health benefits by supporting intestinal homeostasis, maintaining intestinal epithelial integrity, and preventing dysregulation of immune cell activation [20, 21]. Given the established contribution of *A. muciniphila* in maintaining health, oral administration of this bacterium as a probiotic has been investigated in preclinical and clinical studies. Oral gavage of mice with *A. muciniphila* protects against inflammatory bowel disease, obesity, insulin resistance, atherosclerosis, and some cancers [19, 22-25]. In humans, probiotic delivery of *A. muciniphila* was shown to improve metabolic health in overweight or obese individuals [26]. As such, understanding the probiotic effects of this bacterium has important implications for human health. However, very little is known about how *A. muciniphila* colonizes the intestinal tract of mammals in the early stages of life.

Protocols that are able to colonize the gut microbiota of mice with specific bacterial species are needed to understand their effects in preclinical models of disease. Models that orally deliver bacteria are applicable to understanding the probiotic effects of delivered bacteria. The most widely used approach for orally delivering bacteria is oral gavage. However, this method is invasive. It causes distress of animals and can damage the intestinal tract [27, 28]. The procedure is also technically difficult so requires specialized training, especially in studies that use young mice that are physically small. Importantly, differences in the colonization of the gut microbiota in infants leads to long-lasting effects in adults so models are needed to modify the gut microbiota in infant mice [29]. Voluntary feeding of live bacteria has the benefit of being non-invasive, causing less damage, being less technically demanding, and replicating the probiotic delivery of bacteria for preclinical models [30]. It can also be used to examine bacterial delivery to infant mice at early stages of development when they are small but, to our knowledge, has not been extensively studied for this purpose. However, voluntary feeding of mice is challenging because it is difficult to control the timing and amount of material that is ingested. Here, we report the development of a voluntary feeding protocol for delivery of bacteria to colonize the gut microbiota of laboratory mice and use it to examine the colonization of the intestinal tract by *A. muciniphila* early in life. The voluntary feeding protocol is able to deliver a human strain of *A. muciniphila* to the intestinal tract of infant mice during the weaning period whereupon this bacterium stably colonizes the intestinal tract. Using a model of dysbiosis of the gut microbiota that is characterized by severe reduction in *A. muciniphila* [31], we show that voluntary feeding of bacteria in young weaning mice is able to lead to long-term colonization in the gut microbiota. The voluntary feeding protocol effectively modifies an early developmental window in the biology of the gut microbiota and will be a valuable resource for studies on the role of bacterial species in this microbial population. Our findings may also improve the translation of studies on the development of probiotics in preclinical murine models.

## Methods

### Animals & induction of dysbiosis of the gut microbiota

C57Bl/6 mice were purchased from Jackson Laboratories and bred in-house. The gut microbiota was disrupted in mice through administration of an antibiotic cocktail in the drinking water containing ampicillin (0.5 g/L), vancomycin (0.25 g/L), neomycin sulfate (0.5 g/L), and metronidazole (0.5 g/L). Pregnant females were administered antibiotics ad libitum starting before birth and continuing until pups reached 3 weeks of age. Pups were then maintained on regular water thereafter. This treatment prevents colonization of the intestinal tract with bacteria and then permits colonization starting at 3 weeks of age, which is when mice are weaned [32]. The resultant microbiota is similar to conventionally colonized animals but lacks *A. muciniphila* and a handful of other less abundant bacteria [31]. All study protocols were reviewed and approved by the Simon Fraser University Animal Care Committee.

### Culture of *A. muciniphila*

*A. muciniphila* (MucT/BAA-835) was obtained from ATCC. Cultures were grown in a previously described synthetic medium [17], containing 3 mM KH □PO □, 3 mM Na□HPO □, 5.6 mM NH□Cl, 1 mM MgCl□, 1 mM Na□S.9H□O, 47 mM NaHCO□, 1 mM CaCl□, 0.2% GlcNAc, 0.2% glucose, 16 g/L soy peptone (VWR), 4 g/L threonine, and trace elements and vitamins. Cultures were incubated in an anaerobic chamber (Coy Laboratory; 5% H□, 5% CO□, 90% N□) until stationary phase. Cells were harvested by centrifugation, washed once with PBS, and resuspended to approximately 10^1^ □ CFU/mL in sterile PBS containing 20% glycerol. Aliquots were stored at −80°C and thawed immediately before feeding experiments. Thawed bacterial aliquots were serially diluted 2-fold in a 20% glycerol stock in 1xPBS, and OD_600_ was measured. Bacterial recovery was estimated by relating the measured CFU/mL to the expected aliquot concentration, assuming 4.18x10^8^ CFU/mL at OD_600_ = 1.

### Formulation of feeding capsules

Feeding capsules used in all experiments were adapted from Zhang et al [33]. Briefly, a fresh sucralose-gelatin stock was made prior to feeding experiments by dissolving 10% sucralose (wt/vol) and 10% gelatin (wt/vol) in distilled water. Green food coloring was added to the solution and then autoclaved. Autoclaved stocks were aliquoted and stored at 4°C for up to 2 days. Approximately 45 minutes before feeding experiments, the sucralose-gelatin stocks were melted at 37°C for 15 minutes. Glycerol or bacterial aliquots were then thawed at 37°C for 5 minutes and immediately mixed into sucralose-gelatin stock aliquots to the desired bacterial concentration. This mixture was then aliquoted into a 24-well plate (approximately 2g/well) and allowed to solidify for 15 minutes at 4°C. Solidified feeding capsules were cut into 0.25g slices and placed inside animal cages on 6mm petri dishes for consumption.

### Colonization of mice with human *A. muciniphila*

Three days after cessation of antibiotics at 3 weeks of age, mice were administered a 0.25g gelatin capsule containing only sucralose as a training capsule. The day after, mice were given control capsules or capsules containing 10^9^ CFU of human *A. muciniphila* (BAA-835). Mice were fasted for three hours before being fed capsules. Stool samples were collected from mice immediately prior to *A. muciniphila* consumption and at 1, 2, 7, 14 and 28 days after consumption to monitor colonization of the gut by the bacteria.

### Quantification of *A. muciniphila* in fecal samples

DNA was extracted from fecal samples using a QIAGEN QIAamp Fast DNA Stool Mini Kit, and DNA concentration was measured using a nanodrop spectrophotometer. For all PCR and qPCR experiments, 50ng of sample DNA was used. Primers specific to mouse or human strains of *A. muciniphila* were used (Supplemental Figure 1). For PCR experiments, samples were amplified using a Q5 High-Fidelity DNA Polymerase (New England Biolabs) as per kit instructions. PCR products were run on a 1% agarose gel and densitometry was performed on ImageJ. For qPCR experiments, PowerUp SYBR Green Master Mix was used. To quantify human *A. muciniphila* in stool, a five-point DNA standard curve was generated using DNA extracted from a pure culture of BAA-835 with a known CFU/mL.

### Statistical analyses

To determine differences between groups, an ANOVA was performed for comparisons between multiple groups and a Mann Whitney U Test was performed for comparison between two groups.Statistically significant differences were set at p < 0.05.

## Results

### *Akkermansia muciniphila* in the murine gut microbiota

*A. muciniphila* is ubiquitous in the gut microbiota of mammals, although there is variability in its abundance within different hosts because its colonization is affected by diet and competition with other bacteria [34, 35]. Bacteria were quantified in fecal samples from female and male C57Bl/6 laboratory mice using PCR and qPCR that specifically amplifies DNA from *A. muciniphila* strains that are typically found in the murine intestine. *A. muciniphila* was consistently abundant in fecal samples of conventionally housed female and male animals (Fig. 1). Quantitative PCR indicated a potential increase in relative abundance of *A. muciniphila* in males as compared to females (Fig. 1C). Colonization of the gut microbiota with *A. muciniphila* can be affected by environmental conditions, such as antibiotic exposure, and potentially by the time of colonization. We have established that delaying colonization of the female gut microbiota by administering mothers and pups with a broad spectrum of antibiotics until 3 weeks of age leads to dysbiosis of gut microbiota in resultant adults that is characterized by a substantial reduction in *A. muciniphila* [31]. Indeed, early life administration of antibiotics in this way led to a substantial and significant reduction in the abundance of *A. muciniphila* in both female and male mice at 8 – 10 weeks of age (Fig. 1).

**Figure 1.**
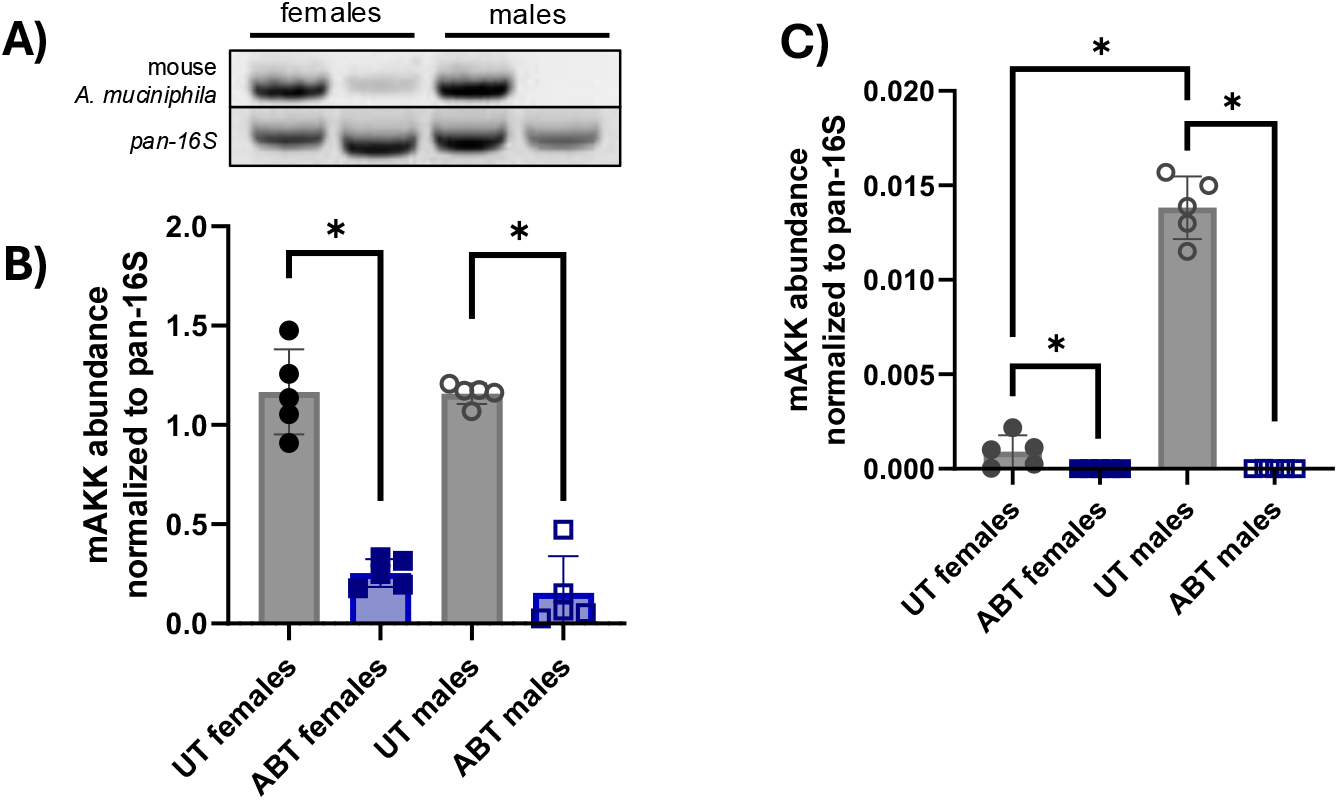
Quantification of *A. muciniphila* in murine fecal samples. A) Representative PCR of mouse *A. muciniphila* (mAKK) in fecal samples from female and male adult mice that were untreated (UT) or given antibiotics until 3 weeks of age (ABT). B) Densitometric analysis of mAKK abundance in UT and ABT female and male mice. C) qPCR analysis of mAKK abundance normalized to *pan-16S* in fecal samples. * p<0.05.

### Voluntary feeding of mice with *A. muciniphila*

Mice can be voluntarily fed material in gelatin capsules containing sucralose, although it is not known whether live bacteria can be delivered in this way and whether this voluntary feeding can be used to colonize the gut microbiota. We developed a voluntary feeding protocol for this purpose (Fig. 2A). Pups were fasted for 3 hours and then gelatin capsules containing only sucralose were prepared and provided to the mice. The recovery of *A. muciniphila* from thawed aliquots was 67% at the time of capsule preparation (Fig. 2B). Introduction of a capsule for the first time led to wariness by mice and slow consumption that was highly variable (Fig 2C). As such, we introduced a training step in which mice were “trained” on the presence of capsules by introducing a control capsule 3 days after weaning (first exposure) and then providing a second “feeding” capsule the day after (second exposure). Introduction of a “training” capsule in this way led to acceptance of the second capsule and full consumption of it within 10 mins by all mice (Fig. 2C).

**Figure 2.**
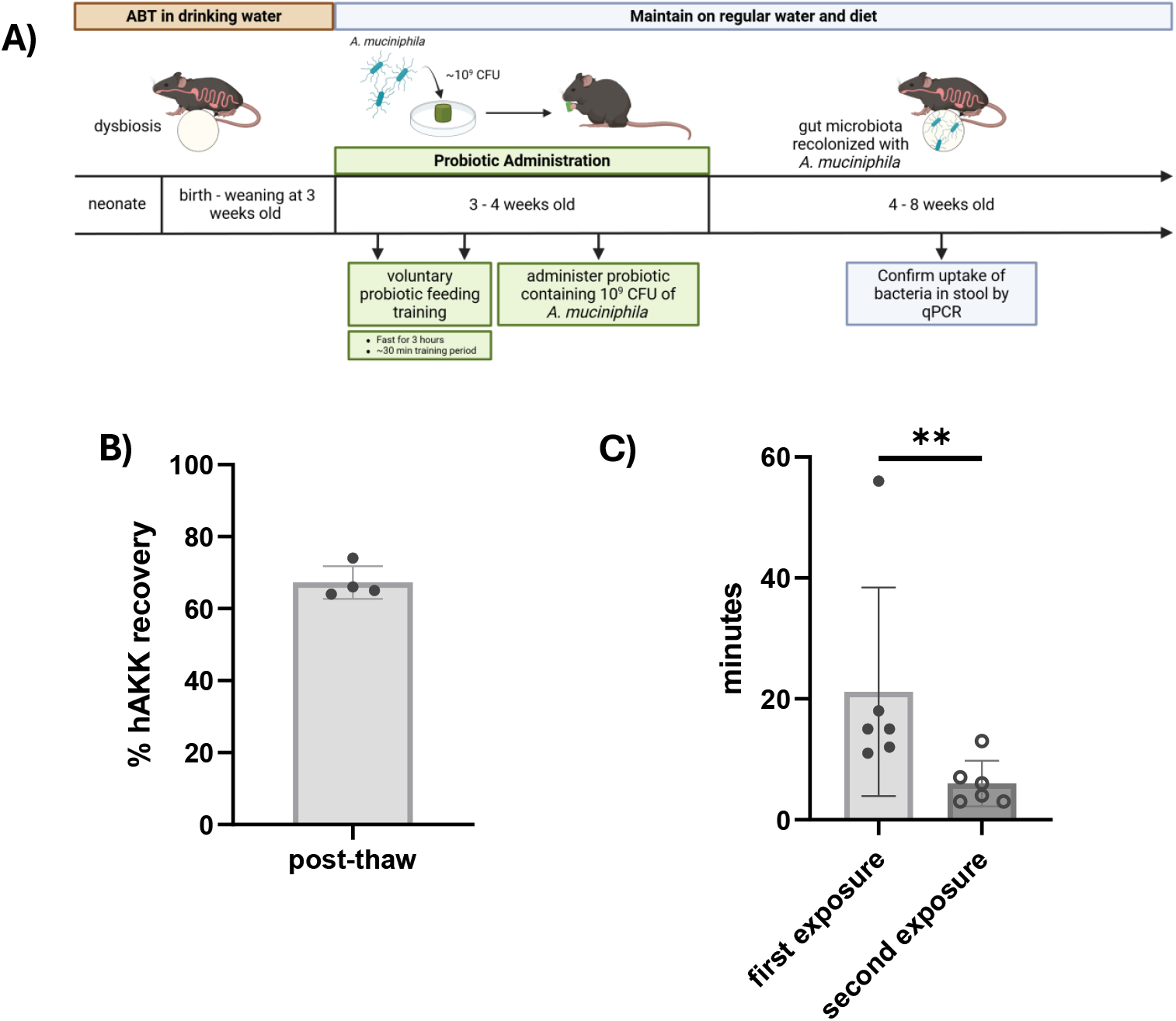
Voluntary oral feeding protocol in mice. A) Schematic of voluntary oral administration protocol. B) Recovery of human *A. muciniphila* (hAKK) immediately after thawing of bacterial aliquots and 45 minutes after embedding in gelatin capsules. C) Consumption time at first and second exposure to gelatin capsules. *p<0.05, **p<0.01

### Colonization of the female gut with *A. muciniphila*

The ability of voluntary feeding to colonize the intestine of female mice with commensal bacteria was then examined by feeding mice *A. muciniphila* (strain BAA-835/MucT; isolated from a human sample [36]. The absolute abundance of human *A. muciniphila* was quantified by qPCR using species-specific primers that detect the human strain, but not the endogenous mouse *Akkermansia*, with quantification based on a standard curve of known bacterial concentrations (Supplemental Figure 2).Female mice were then fed control capsules or capsules containing human *A. muciniphila* and fecal samples analyzed. Human *A. muciniphila* was not detected in mice that were fed control capsules. In mice that received the bacterium, human *A. muciniphila* was abundant in fecal samples as early as day 1 after feeding and its abundance was maintained until at least 28 days, which was the last time-point studied. Abundance of this bacterium in fecal samples was stably maintained at 10^5^ – 10^6^ CFU pergram of stool, although there was variability at specific time-points (Fig. 3A). Feeding mice *A. muciniphila* did not affect the weight of mice (Fig. 3B).

**Figure 3.**
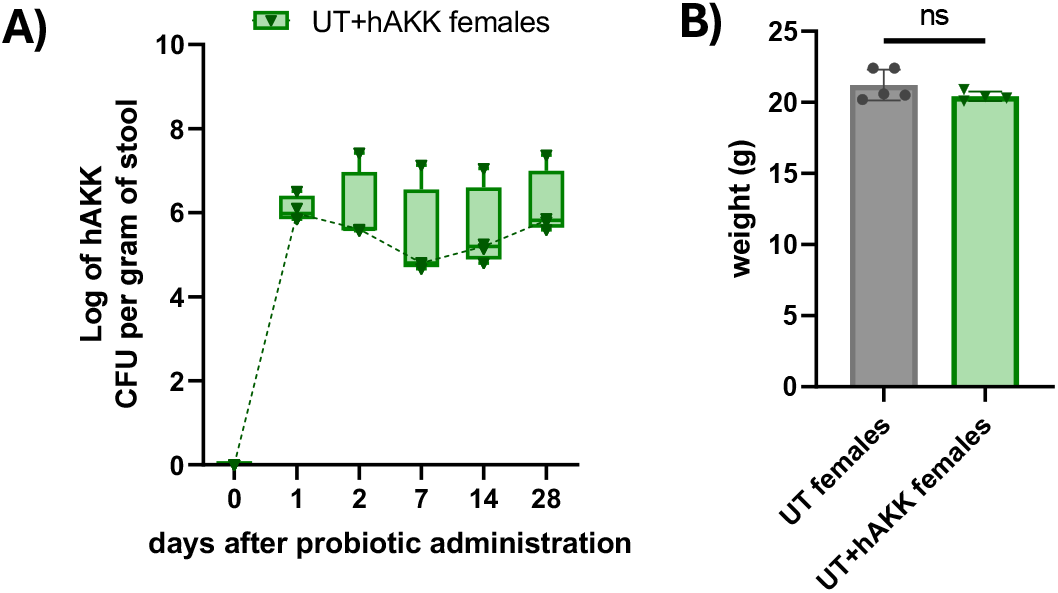
Enhancement of Akk in UT female mice. A) Absolute quantification of human *A. muciniphila* (hAKK) in female fecal samples. B) Mouse weights.

In addition to examining the colonization of an undisturbed gut microbiota, we also evaluated how voluntary feeding can colonize dysbiotic gut microbiota in which the presence of bacteria was prevented by broad spectrum antibiotics until 3 weeks of age. This is a developmental window at which weaning occurs and is an inflexion point in the development of the gut microbiota [37]. Female mice and their mothers were maintained on water containing broad spectrum antibiotics that reduces the total amount of bacteria in the intestinal tract by greater than 99% [32]. At 3 weeks of age, antibiotics were stopped and mice were maintained on normal water. Mice were then given a training capsule 3 days after the cessation of antibiotics and then given a control capsule or capsule containing human *A. muciniphila* 1 day later. The abundance of endogenous mouse *A. muciniphila* was significantly reduced by treatment with antibiotics and feeding with human *A. muciniphila* did not affect this (Fig. 4A-C). Human *A. muciniphila* was absent from the gut microbiota of control mice and mice that were given antibiotics (Fig. 4A-C). When mice were fed human *A. muciniphila*, the strain was detectable in fecal samples beginning at day 1 after consumption and lasting until the last measurement at 28 days after consumption (Fig. 4D). Colonization of the gut microbiota was rapid, with variability at early time-points after feeding and then stabilization with less variability between mice after day 7 post-consumption. The abundance of human *A. muciniphila* was 10^7^ – 10^8^ CFU per gram of feces, which was higher than in mice not given antibiotics (Fig. 4D). The weights of mice were not affected by feeding with human *A. muciniphila* (Fig. 4E).

**Figure 4.**
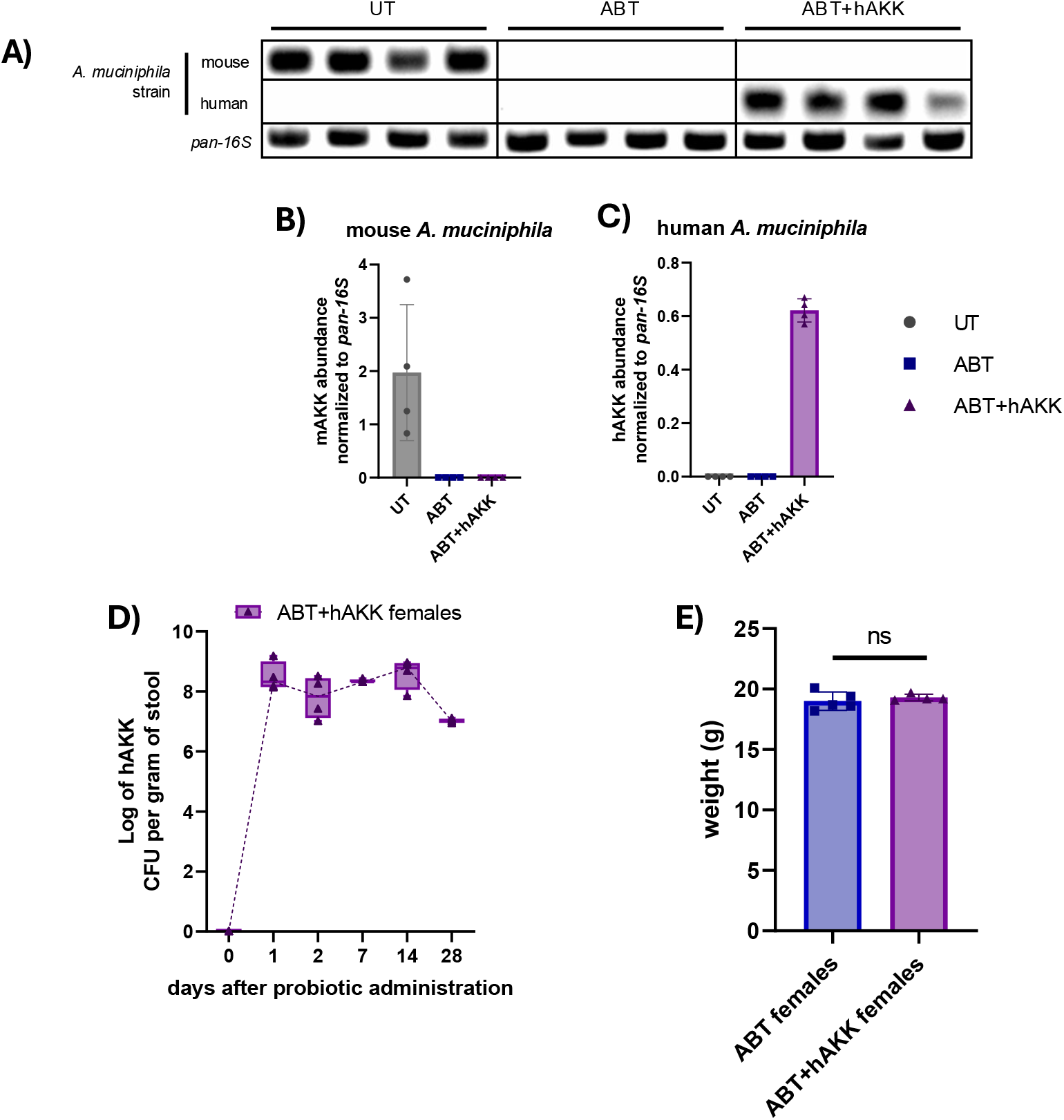
Colonization of ABT mice with human *A. muciniphila*. A) PCR of human *A. muciniphila* (hAKK) and mouse *A. muciniphila* (mAKK) DNA in the stool of UT, ABT and ABT + hAKK female mice. Densitometric analysis of B) mAKK and C) hAKK abundance normalized to 16S in fecal samples. D) Absolute quantification of hAKK in ABT + hAKK female stool samples by qPCR. E) Mouse weights.

### Colonization of the male gut microbiota with *A. muciniphila*

Colonization of male mice with human *A. muciniphila* was also examined. Human *A. muciniphila* was not detected in fecal samples from male mice at the time of weaning. In untreated male mice, human *A. muciniphila* was abundant as early as day 1 after feeding and levels were maintained until at least day 28 after feeding, which was the last time-point tested. There was a transient reduction in bacterial abundance at day 7 after feeding that rebounded by day 14. With the exception of day 7, *A. muciniphila* levels were maintained at an abundance of 10^6^ – 10^7^ CFU per gram of stool at all time-points (Fig. 5A).

**Figure 5.**
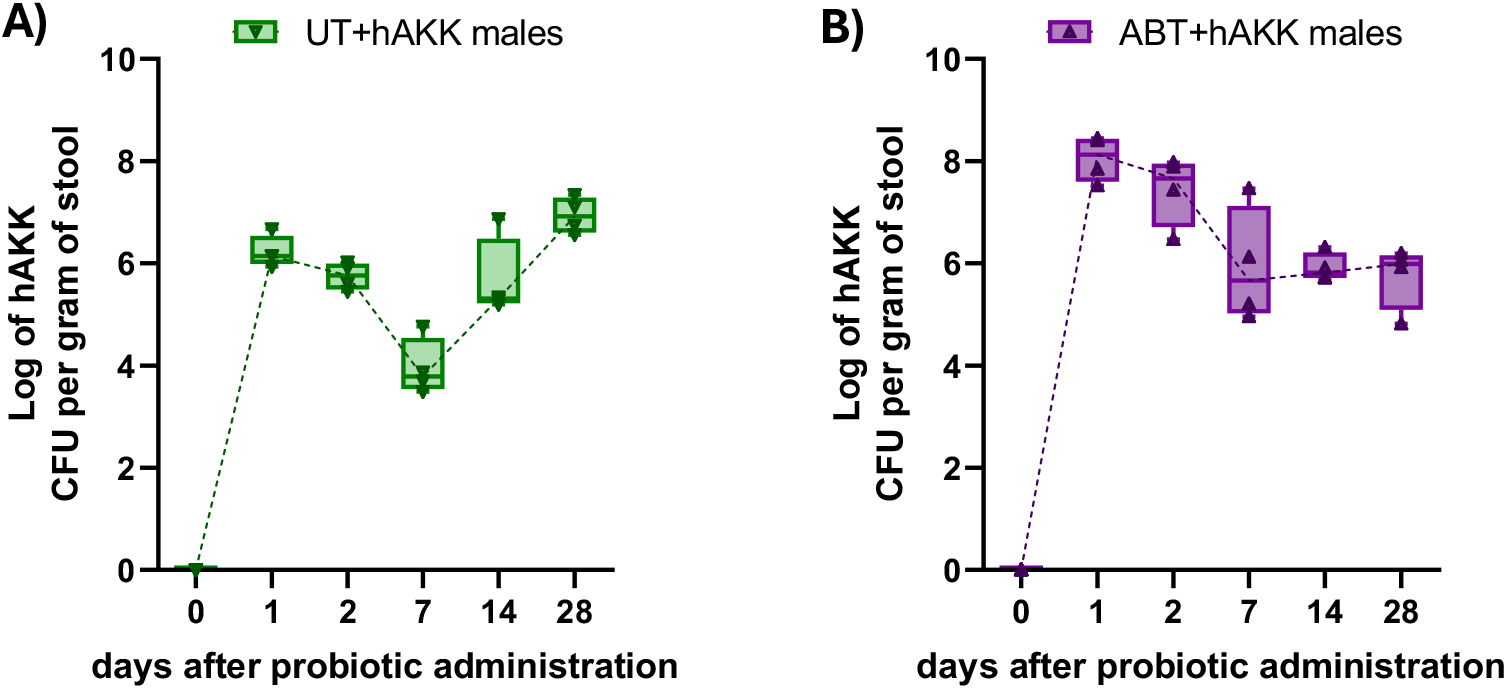
Colonization of male mice with human *A. muciniphila*. Absolute quantification of human *A. muciniphila* (hAKK) in A) UT + hAKK and B) ABT + hAKK male fecal samples over time.

Colonization of male mice that received antibiotics for the first 3 weeks of life was also examined. In these mice, human *A. muciniphila* was abundant at 10^8^ CFU per gram of stool in fecal samples as early as day 1 after feeding. There was a subsequent decline in abundance to 10^6^ CFU pergram of stool at day 7 after feeding and then bacterial levels were maintained at this abundance until at least day 28 after feeding (Fig. 5B).

#### *A. muciniphila* incompletely occupies its niche in the week after weaning

Because human *A. muciniphila* colonized the intestinal tract in the absence of pretreatment of mice with antibiotics, we investigated whether this bacterium incompletely occupies its niche in the week after weaning and whether this could lead to its competitive removal by another strain of *A. muciniphila* during this developmental window. When the abundance of mouse *A. muciniphila* (that naturally colonizes the intestinal tract starting shortly after birth) was quantified, this bacterium was present in fecal samples at the beginning of the weaning period (week 3 of life). The relative abundance of this bacterium was significantly increased in adult males (7 – 8 weeks of life) and numerically increased in adult females, although the latter did not reach statistical significance (Fig. 6). When mice were fed human *A. muciniphila*, the endogenous mouse *A. muciniphila* was eliminated in adult mice (Fig. 6). The elimination of mouse *A. muciniphila* coincided with the colonization by human *A. muciniphila* after voluntary feeding. This finding indicates that voluntary feeding with human *A. muciniphila* competitively excluded the mouse strain from colonization of the intestinal tract.

**Figure 6.**
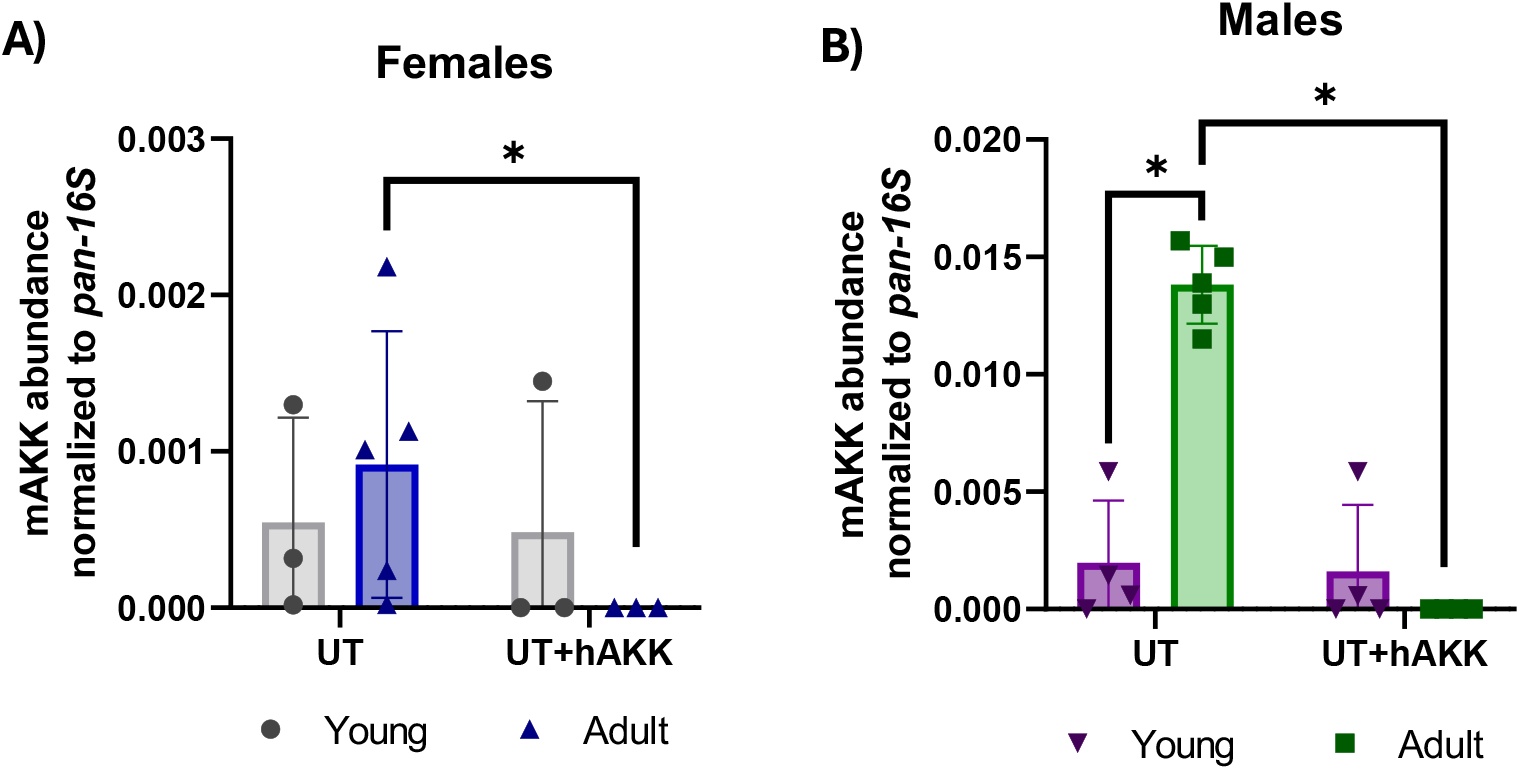
Colonization of mice with endogenous mouse *A. muciniphila* after weaning. Quantitative PCR of endogenous mouse *A. muciniphila* (mAKK) DNA normalized to the 16S rRNA gene in mouse stool DNA. Stools were collected from young mice immediately after weaning (3 weeks of age), then in adulthood (7-10 weeks of age) from control mice that did not receive treatment (UT) or that received human *A. muciniphila* (UT + hAKK). *p<0.05.

## Discussion

We have established a voluntary feeding protocol that is able to colonize the intestinal tract of mice with the gut commensal *A. muciniphila*. The protocol rapidly colonizes female and male mice in a reproducible manner and can be applied to animals that are housed normally or those in which colonization by the gut microbiota is delayed. Voluntary feeding mice in this way enabled the characterization of intestinal colonization by *A. muciniphila* early in life, showing that this bacterium rapidly colonizes the intestinal tract when delivered shortly after weaning. The ability to colonize mice with gut bacteria using this non-invasive method will be a benefit for studying the gut microbiota, especially in young mice, and for researching probiotics in murine models.

Oral gavage remains the main method to introduce microbes into the intestinal tract of mice. However, it is inherently invasive. Studies have reported damage to the digestive tract of mice after gavage and the technical demands can lead to variations in the amount of bacteria delivered into the stomach [27, 38]. Oral gavage is also difficult to perform in young mice that are small, which is an important developmental age in which the gut microbiota is developing and malleable. We show that voluntary feeding of young mice with sucralose-sweetened gelatin capsules is able to deliver live bacteria into the intestinal tract of 3 – 4 week old mice. Mice were inherently wary of the capsules upon the first exposure and a training capsule was needed in order to enable the fast and consistent consumption of a second capsule that contained bacteria. The protocol was non-invasive and mice did not show signs of distress. Also, the protocol did not affect the resultant weight of adult mice.

The timing of bacterial delivery is likely to affect its colonization in the intestinal tract. The gut microbiota is malleable during infancy and after weaning [37]. Indeed, the physiological response to intestinal colonization with microbes during weaning is essential for defining lifelong homeostasis of the immune system [39]. *A. muciniphila* begins colonizing the intestinal tract before weaning due to its ability to degrade oligosaccharides in breast milk [40]. As such, environmental exposures have a considerable influence on how the gut microbiota colonizes during these early stages of life and disrupting healthy colonization until after weaning leads to establishment of a dysbiotic population in adults that has pathogenic implications [41]. Notably, exposure of humans to common antibiotics around the time of weaning reduces *A. muciniphila* in their gut microbiota while similar exposures in adults does not affect this bacterium [42-44]. This may have health implications because antibiotic exposure in infancy and reduced *A. muciniphila* are associated with many diseases [11]. Voluntary feeding of young mice enables the modification of the gut microbiota to examine the effects and therapeutic benefits of specific bacterial species that is applicable to many preclinical models. It is unclear if voluntary feeding in adults can lead to colonization. In studies that use oral gavage to colonize adult mice with *A. muciniphila*, treatment with antibiotics prior to delivery is needed to create niches for the delivered bacteria to thrive [45].

We observed that colonization with *A. muciniphila* was rapid when mice were fed between 3 – 4 weeks of age. The delivered bacterium was abundant in fecal samples by day 1 after feeding and was stably maintained until 28 days after feeding, which was the latest time-point examined. The persistence of *A. muciniphila* in stool samples after a single feeding is indicative of intestinal colonization with live bacteria. The kinetics of colonization were the same in normally colonized female and male mice, regardless of whether mice were maintained on antibiotics or not prior to feeding. Colonization of the intestinal tract with *A. muciniphila* is governed by competitive exclusion whereupon the presence of an existing strain prevents colonization with a separate one [46]. We observe that human *A. muciniphila* colonized the intestinal tract despite the endogenous presence of mouse *A. muciniphila* at the time of feeding. This may occur because the abundance of mouse *A. muciniphila* was low at the time of feeding, leading to incomplete occupation of its niche. Indeed, exposure to antibiotics resulted in higher concentrations of *A. muciniphila* in both sexes early after feeding, which is consistent with a lack of competition with other bacteria for mucin niches in mice that received antibiotics. The stabilization of bacterium to similar levels in all mice suggests that the size of the *A. muciniphila* population is endogenously defined by its niche. Interestingly, the oral gavage of 5 week old mice (that were treated with tetracycline) with *A. muciniphila* leads to intestinal colonization at 10^2^ – 10^3^ greater levels than our voluntary feeding protocol [45]. Thus, our findings show that colonization of the mouse intestinal tract with *A. muciniphila* is active shortly after weaning but remains malleable because it incompletely occupies its niche until at least 4 weeks of life. This is consistent with the kinetics of its colonization of the human gut microbiota [40].

Several studies have reported differences in the abundance and effects of *A. muciniphila* in females and males [47]. In clinical studies, the relative abundance of *A. muciniphila* may be higher in females that in males [48]. Interestingly, we observed a relatively higher abundance of *A. muciniphila* in the endogenous gut microbiota of male as compared to female mice. The abundance and effects of *A. muciniphila* are differentially affected by diet in females when compared to males [49, 50]. Also, the diversity of the gut microbiota may be lower in males than females, which may affect the relative abundance of specific species [51]. There may be sex-dependent effects of *A. muciniphila* on immune responses because exacerbation of immune-mediated transplant vascular injury is exacerbated by dysbiosis in female, but not male, graft recipients that is characterized by reduced *A. muciniphila* [31]. We find that voluntary feeding of mice with *A. muciniphila* was able to similarly colonize female and male mice, although there may be distinctions in the kinetics of colonization. In untreated mice of both sexes, colonization occurred very quickly (one day) at apparently similar levels. There may be a transient reduction of bacterium at day 7 after feeding in male mice that rebounded to stable levels that were comparable to those in females. In antibiotic-treated mice, *A. muciniphila* colonized the intestinal tract of female and male mice with similar kinetics.

Voluntary feeding of bacteria can model the delivery of microbes to the intestinal tract as probiotics. The abundance of *A. muciniphila* is associated with many aspects of human health so strategies to increase its colonization of the intestinal tract may have health benefits [52, 53]. Indeed, *A. muciniphila* is extensively studied as a potential probiotic for several conditions. Preclinical studies using oral gavage show that its delivery in this way reduces obesity, metabolic syndrome, inflammatory bowel disease, and atherosclerosis [23, 25, 54]. Studies in adult mice show that some of these therapeutic effects may not require colonization by live bacterium and are mediated by cell surface components of this microbe that signal to intestinal cells [17]. Further, clinical trials have shown that probiotic delivery of *A. muciniphila* reduces metabolic abnormalities [55]. This bacterium is considered a strict anaerobe although it can survive for several hours – days in aerobic conditions [56, 57]. As such, translating its use as a probiotic will need to consider this in the preparations.

## Conclusion

We have developed a voluntary feeding protocol for colonizing the intestinal tract of mice with commensal bacteria and used it to characterize the early life colonization by *A. muciniphila*. The protocol has implications for preclinical development of probiotics and understanding the biology of commensal bacteria. Given the increasing role of *A. muciniphila* as a probiotic for the potential treatment of several human diseases, understanding its colonization and delivery into the intestinal tract may have health implications.

## List of abbreviations

CFU: colony forming units
PCR: polymerase chain reaction
SCFA: short-chain fatty acid

## Declarations

### Ethics approval and consent to participate

All studies were reviewed and approved by the Simon Fraser University Animal Care Committee.

### Consent for publication

All authors consent to publication of the data.

### Availability of data and materials

The data in this study are available from the corresponding author upon reasonable request.

### Competing interests

There are no financial or non-financial competing interests to report.

### Funding

The work was funded by a Grant-in-Aid from the Heart and Stroke Foundation of Canada (J.C.C) and a Discovery Grant from the Natural Sciences and Engineering Research Council of Canada (L.D.)

## Authors’ contributions

JMD generated data, analyzed data, and wrote the manuscript; PM generated data and analyzed data; CH generated essential materials for the study; EG generated data and analyzed data; WE generated data and analyzed data; LD provided essential materials for the study, analyzed data, and wrote the manuscript; JCC analyzed data and wrote the manuscript.

## Notes

### Competing Interest Statement

The authors have declared no competing interest.

