## Supplemental figures for "Rapid early life colonization of the intestinal tract by *Akkermansia muciniphila* after voluntary feeding"

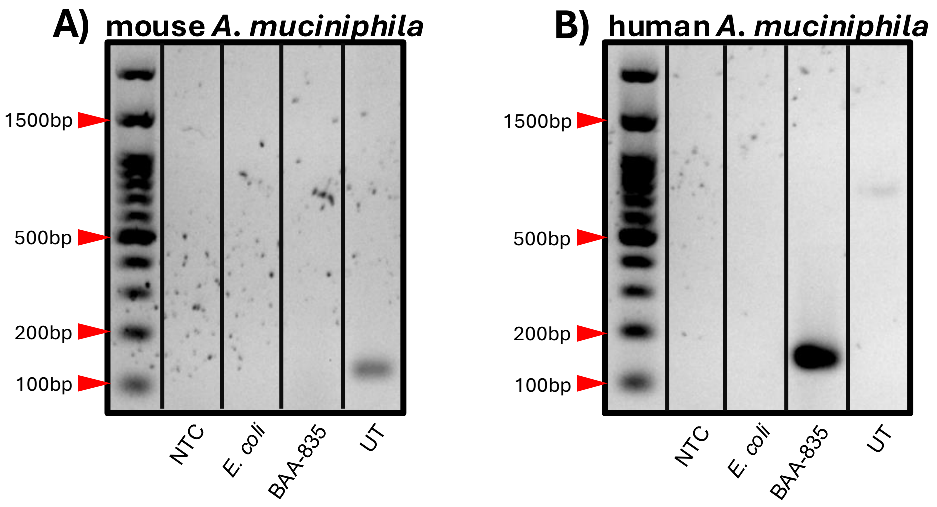


C)

| Target | Sequence |
|---|---|
| mouse<br>A. muciniphila | Forward: 5’-TGATTCCGATACGGACGCTG-3’<br>Reverse: 5’-TTGCCATTTCCCCGGTACTC-3’ |
| human<br>A. muciniphila | Forward: 5’-GCGGCTTGCTCAAAACATTG-3’<br>Reverse: 5’-TGCCAAAAGTGCCCTTTCTG-3’ |

**Figure S1. Strain specific *A. muciniphila* primers.** PCR using mouse and human *A. muciniphila* primers. To check for specificity, DNA from a non-pathogenic *E.* coli, a pure culture of human *A. muciniphila* (MucT/BAA-835) and from untreated mouse stools (UT) were used. A negative control template (NTC) containing no DNA was used as a loading control. A) mouse *A. muciniphila* specificity. B) human *A. muciniphila* specificity. C) Strain specific *A. muciniphila* primer sequences.

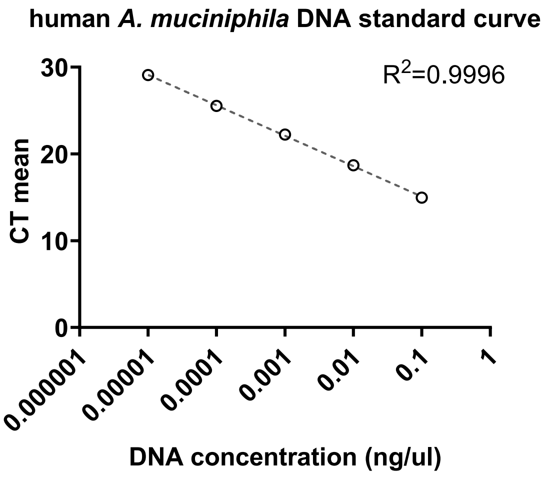


**Figure S2. qPCR DNA standard curve of human *A. muciniphila*.** A 5-point DNA standard curve was generated using DNA from a pure human *A. muciniphila* bacterial aliquot of known CFU/mL.
